# Doxorubicin Induces Senescence in Intestinal Epithelial Cells

**DOI:** 10.1101/2021.01.29.428764

**Authors:** Mandy Biraud, Jocsa Cortes, Paul Cray, Guy Kunzmann, Javid Mohammed, Christopher M. Dekaney

**Author notes:** Corresponding author, Address: 1060 William Moore Drive, Campus Box 8401, Raleigh, North Carolina 27607.

## Abstract

Doxorubicin treatment induces DNA damage and apoptosis in rapidly dividing cell types like intestinal epithelial cells. This has been demonstrated both *in vivo* and *in vitro*. In certain cell types some cells do not undergo DNA damage-induced apoptosis in response to doxorubicin but instead become senescent. Induction of senescence in these cells can lead to dysfunction and chronic inflammation, which can lead to more damage. We questioned whether a single dose of doxorubicin would be able to induce apoptosis and senescence in intestinal epithelial cells *in vitro*. For these studies, we exposed IEC-6 small intestinal epithelial cells to doxorubicin to evaluate whether senescence is induced in a relatively homogeneous population of intestinal epithelial cells. Although some cells underwent apoptosis, those that did not showed traits of senescence. Our studies showed that doxorubicin treatment increased cell size and increased expression of senescence-associated β-galactosidase. Concomitantly, we observed increased mRNA expression of several genes associated with a senescence-associated secretory phenotype including *IL-6*, *Ptges*, *Faim2*, and *Cdkn1a* and decreased expression of *Sirt1*. We also observed release of HMGB1, a cellular alarmin, from treated cells. Together, these data suggest that doxorubicin induces senescence in intestinal epithelial cells. Furthermore, our data indicate that cellular responses to a DNA damaging agent, such as doxorubicin, can differ within a population of cells suggesting differing levels of sensitivity within a relatively homogenous cell population. Further studies are needed to delineate the mechanisms that determine whether a cell moves down an apoptotic or senescent pathway following DNA damage.

## Introduction

Doxorubicin (DXR) is a chemotherapeutic widely used to treat a variety of cancers including breast, lung, and ovarian cancers. Off target effects of DXR are manifested in the development of mucositis, diarrhea, and vomiting [0]. Using a murine model, we have demonstrated that DXR administration leads to a sequela of damage to the intestinal mucosa; rapid induction of DNA damage and apoptosis within 6 hours of treatment followed by cell cycle arrest is observed in the crypt epithelium.(1) Despite this change in number of and proliferative status of cells within intestinal crypts, evidence of significant intestinal crypt loss is not observed until 72 hours after DXR.(1) The mechanisms responsible for this gradual loss of intestinal crypts is not well understood. In other cell types, including cardiomyocytes, agents such as DXR induce DNA damage and senescence (2,3). This led us to consider whether DXR treatment could induce senescence in intestinal epithelial cells.

Senescence is a cellular process by which the cell terminally exits the cell cycle, and experiences what is termed the senescence associated secretory phenotype (SASP).(4) SASP is characterized by the upregulation and secretion of chemokines and cytokines including IL-6, CXCL1, FAIM2, and PTGES.(4) Although cells are no longer cycling they remain metabolically active and are distinct from quiescent cells. Depending upon the context, induction of senescence in response to DNA damage can have positive and negative impacts within a tissue. For example, induction of senescence within cancerous tissue prevents tumors from expanding.(5) In contrast, induction of senescence in normal tissues can cause early aging and accumulation of senescent cells which have been shown to lead to an increased accumulation of local inflammatory cells.(6,7)

While cellular senescence has been shown by others to result from DXR treatment in other cell types and has been studied in the context of aging of the small intestinal epithelium, we do not know whether DXR induces senescence in intestinal epithelial cells. We hypothesized that DXR treatment would induce cellular senescence in intestinal epithelial cells as defined by changes in cellular morphology and expression of senescence associated genes. We chose to use the IEC-6 cell line as they are a relatively homogenous proliferating population of cells. Our data demonstrate that while DXR does rapidly induce an increase in cleaved caspase 3 expression in some cells other cells take on a senescent phenotype as evidenced by increased expression of *Cdkn1a*, upregulation of genes associated with SASP, increased expression of SA-β-gal, and increased cell size.

## Materials and methods

### Cell culture

Normal rat intestinal epithelial IEC-6 cells were obtained from the American Type Culture Collection (#CRL-1592, Manassas, VA) and grown in in Dulbecco’s Modified Eagle’s Medium (DMEM) supplemented with 5% fetal bovine serum (FBS; Gemini, Sacramento), 2 mM glutamine, and 0.1 U/μmL insulin in a 5% CO_2_ incubator at 37° C. Cells were maintained by changing the medium twice a week and passaged using 0.05% trypsin/ EDTA. For assays, cells were cultured in 6- (Olympus) and 96-well plates (Falcon) at a density of 2 × 10^4^ and 1 × 10^4^ cells per well, respectively and were used within 10 passages. Mycoplasma testing was conducted (LookOut Mycoplasma qPCR Detection Kit, Sigma-Aldrich, Saint-Louis, MO) and found to be negative.

### MTT assay

To measure cell viability, IEC-6 cells were seeded on a 96-well plate. Two days after seeding, cells were treated with media containing 0.2, 0.4, 0.8, 1, or 2 μg/mL of doxorubicin hydrochloride (DXR) (Actavis Pharma, Parsippany-Troy Hills, NJ) for 24 hours or 4 days. After treatment, cell viability was measured using the Vybrant® MTT Proliferation Assay kit (V13154 Thermo Fisher Scientific, Waltham, MA) following the manufacturer’s instructions.

### Senescence associated beta-galactosidase staining

IEC-6 cells were plated in a 6-well plate for two days before DXR treatment (0.2 μg/mL). After incubation with DXR for 4 days, β-galactosidase activity was detected using the Senescence β-Galactosidase Staining Kit (#9860S, Cell signaling Technology, Danvers, MA) according to the manufacturer’s instructions. Briefly, the cells were washed once with 1X PBS and then placed in fixative solution at room temperature for 15 min. Cells were rinsed twice with 1X PBS and stained with β-Galactosidase Staining Solution at 37°C in a dry incubator (no CO_2_) overnight. Cells were observed under a light microscope for the presence of blue precipitate. The presence of blue granules within the cytoplasm of cells were indicative of a positive result for β-Galactosidase staining.

### Cellular senescence live cell analysis assay by flow cytometry

Analysis of cellular senescence was carried out using a Cellular Senescence Live Cell Analysis Assay Kit (ENZ-KIT130-0010# Enzo Life Sciences, Farmingdale, NY). IEC-6 cells were plated in a 6-well plate for two days before DXR treatment. After 4 days of DXR treatment, cells were placed in pretreatment solution at 37°C for 2 hours and stained with SA-β-Gal substrate at 37°C for 4 hours. Cells were washed three times with 1X PBS, trypsinized, washed in 2% FBS/1 X PBS and analyzed via flow cytometry (Becton Dickinson LSRII, Franklin Lakes, NJ). A 488nm Blue laser was used as an excitation source, and the emission was captured using 505LP (505nm Long Pass) and 530/30nm Band Pass optical filters. Data was acquired using BD FACS Diva software, and the histogram-overlays were obtained using FlowJo software.

### IEC-6 cell area measurements

Five brightfield images were taken per well at 10x magnification. A total of 6 wells per treatment were used for these measurements. A minimum of 30 cells per image were measured. Each image was normalized to enhance contrast to show the cellular body using Fiji, and images were measured in a blinded fashion. Cells were outlined by hand and cell area was calculated using Fiji. Super plot was generated using the seaborn python package.

### RNA isolation and quantitative RT-PCR

Total RNA was isolated using NucleoSpin TriPrep (Macherey Nagel, Bethlehem, PA) following the manufacturer’s instructions. cDNA was generated utilizing Applied Biosystems High Capacity Reverse Transcription cDNA kit Applied Biosystems (Waltham, MA), Quantitative real-time PCR was performed in triplicates using the QuantStudio 6 Flex Real-Time PCR System and Taqman Universal PCR Master Mix (Applied Biosystems, Foster City, CA). Probe sets for *Il6* (Rn01410330_m1), *Ptges* (Rn00572047_m1), *Faim2* (Rn01532650_m1), *Cxcl1* (Rn00578225_m1), *Hmgb1* (Rn02377062_g1), *Ascl2* (Rn00580387_m1), *Lgr5* (Rn01509662_m1), *Cdkn1a* (Rn00589996_m1), *Sirt1* (Rn01428096_m1), *Rnf43* (**Rn01471897_m1**), *Myc* (**Rn07310910_m1**), *Axin2* (**Rn00577441_m1**), *Olfm4* (**Rn01465807_m1**), and *β-actin* (Rn00667869_m1) were purchased from Applied Biosystems. Data was analyzed using the ΔΔC_t_ method with normalization to *β-actin* mRNA of the untreated IEC-6 cell group.

### Immunoblotting analysis

IEC-6 cells cultured in 6-well plates were incubated with 0.2 μg/mL of DXR for 4 days. The cells were washed with PBS and lysed with ice-cold radioimmunoprecipitation assay (RIPA) buffer containing 200 mM PMSF, protease inhibitor cocktail and 100 mM sodium orthovanadate (Santa Cruz Biotechnology, Dallas, TX). Protein concentration was measured using the BCA Protein Assay Kit (Thermo Fisher Scientific, Waltham, MA) following the manufacturer’s instructions. Samples were mixed with an equal volume of Laemmli Buffer (Biorad, Hercules, CA) and PBS, and boiled for 5 min. Samples were separated by SDS electrophoresis through polyacrylamide gels (Biorad, Hercules, CA) and transferred to PVDF membranes using the Trans-Blot Turbo Mini PVDF Transfer Kit (Biorad, Hercules, CA). Membranes were blocked with 5% milk in Tris-buffered saline containing 0.1% Tween 20. Membrane was incubated with primary antibodies HMGB1 (0.5 μg/mL, #PA1-16926, Thermo Fisher Scientific, Waltham, MA), and β-actin (1/2000, #4970, Cell Signaling Technology, Danvers, MA) in 5% milk in PBS overnight at 4°C. Membrane was washed and incubated with secondary goat anti-rabbit horseradish peroxidase linked secondary antibody (1/2000, #ab6721, Abcam, Cambridge, MA) for 1 hour. HRP signal was detected using the Clarity Western ECL Substrate (Biorad, Hercules, CA).

### Statistics

All results are presented as means ± SD. Figure legends indicate the respective sample sizes used per experiment. GraphPad Prism 8.0 (GraphPad Software, San Diego, CA) was utilized for statistical analysis. Analysis involving the comparison of two groups were analyzed via Welch’s t-test, and those with more than two groups were analyzed via one-way ANOVA. Statistical significance was determined when *P* <0.05.

## Results

### Dose concentration effect in DXR-induced cell viability

We performed a dose response study with various doses of DXR to determine the optimal dose of doxorubicin (DXR) to use to study the induction of senescence while not completely reducing cell viability. IEC-6 cells were grown in complete DMEM media in the presence of vehicle (control) or presence of different concentrations of DXR for 4 days and cell viability was measured by MTT assay. DXR significantly decreased IEC-6 cell viability in a dose dependent manner 4 day after treatment (Figure 1). The lowest concentration of DXR (0.2 μg/mL) significantly decreased overall cell viability compared to the untreated control at 4 days. Overall, the lowest concentration of DXR (0.2 μg/mL) was chosen for the remaining experiments in this study.

**Figure 1.**
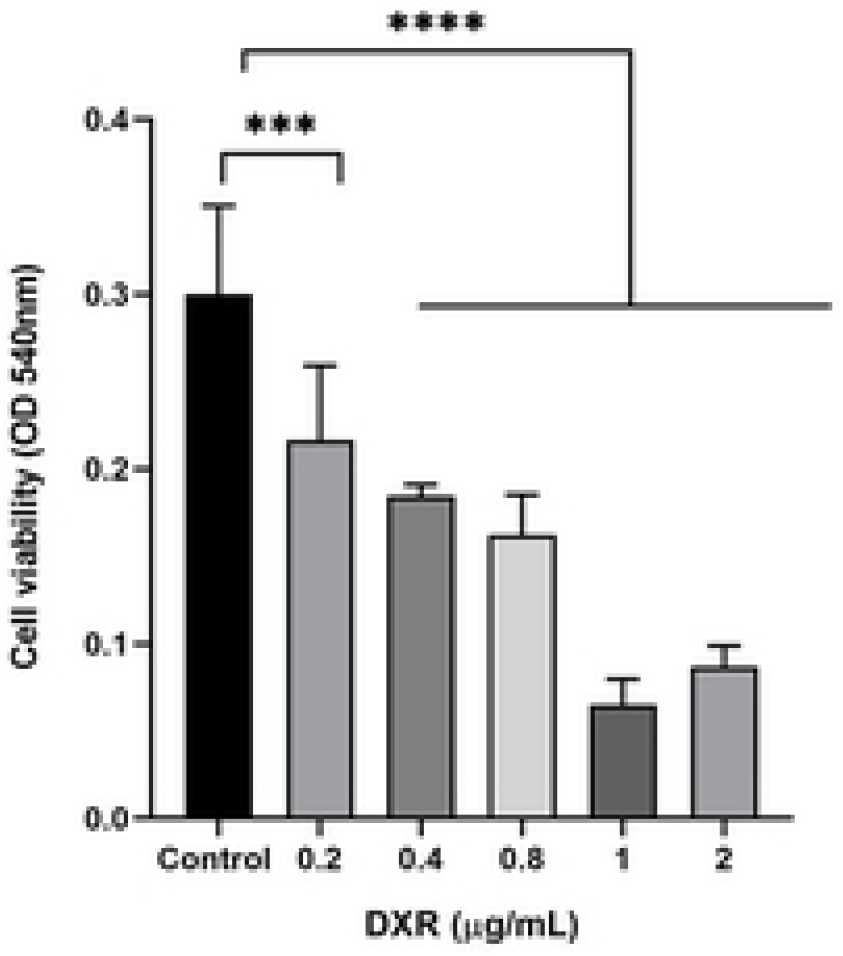
DXR treatment reduces IEC-6 cell viability in a dose-dependent manner. IEC-6 cells were cultured in the presence of vehicle or 0.2, 0.4, 0.8, 1 or 2 μg/ml DXR for 4 days and cell viability was assessed by MTT assay as described in materials and methods. The data are expressed as means ± SD of absorbance values from 3-6 separate wells *** P < 0.001, **** P<0.0001.

### DXR-induced apoptosis is time dependent

Upregulation of cleaved caspase-3 (CC3) protein is a hallmark of cellular apoptosis, and therefore used as surrogate to quantify levels of apoptosis at various durations of DXR treatment. CC3 protein in cells was detected via immunofluorescence staining in vehicle treated cells (Control) and cells treated with 0.2 μg.ml DXR for 6, 12, 18, or 24 hours. Immunofluorescence staining demonstrate an increase in CC3-positive cells at 12 and 24 hours after DXR treatment (Figure 2A). Quantification of the percentage of positive cleaved caspase-3 cells per field demonstrated significantly higher percentages at 12, 18, and 24 hour treatment groups compared to control cells with a maximal percentage of 15% at 24 hours (Figure 2B). These data suggest that at this dose of DXR not all cells undergo apoptosis.

**Figure 2.**
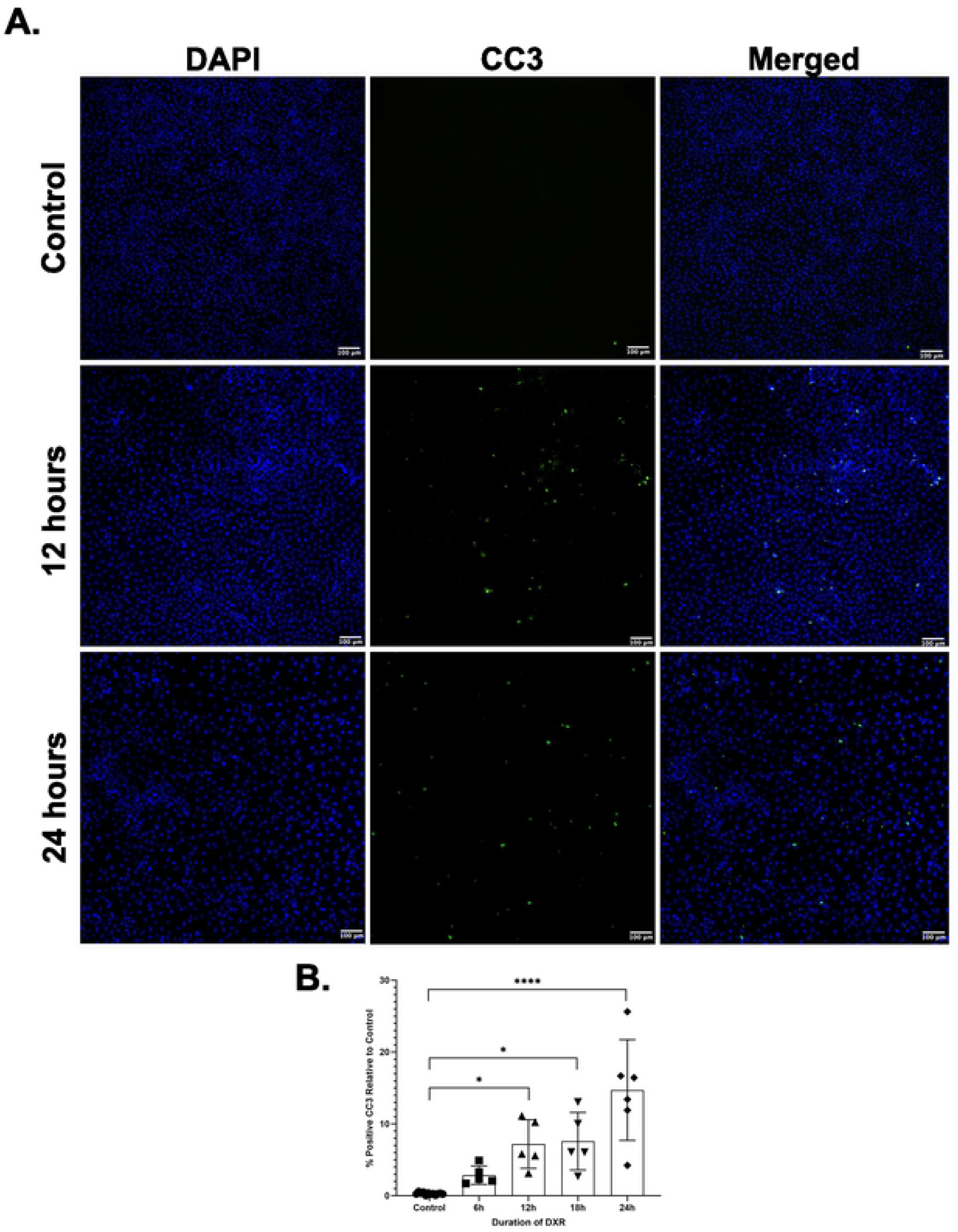
The percentage of cleaved caspase-3 positive cells increases with increasing duration of DXR treatment. Representative immunofluorescence images of IEC-6 cells stained for cleaved caspase-3 and DAPI; untreated control, 12h and 24h post-treatment with 0.2μg/ml. Scale bar equals 100 μm. (B) IEC-6 cells were treated with DXR (0.2μg/ml) for 6-24 hours and positive cleaved caspase-3 cells expressed as percentages of DAPI positive cells. n= 5-6 for each condition, error bars represent the standard deviation (SD). Significance was calculated using the Kruskal-Wallis’s test, * P<0.05, **** P<0.0001.

### DXR treatment increases cell size

Increase in cell size is observed in other cells types entering cellular senescence. Therefore, we treated IEC-6 cells with vehicle or 0.2 μg/ml DXR for 4 days and then brightfield images were collected for analysis of cell area. As can be seen in Figure 3A, DXR treatment increased the size of IEC-6 cells compared to vehicle treated controls. Quantification of cell area (μm^2^) demonstrated the significant increase in cell size associated with DXR treatment which resulted in an almost 10-fold increase in cell area (Figure 3B). Cells were then trypsinized to individual cells and subjected to flow cytometry to measure forward scatter, a surrogate measurement of cell size. As can be seen in Figure 3C, DXR treatment resulted in substantial increases in both side scatter (measurement of cellular granularity) and forward scatter. Quantification of average cellular forward scatter showed a significant increase in average cellular forward scatter values for DXR treated cells compared to vehicle treated cells (Figure 3D).

**Figure 3.**
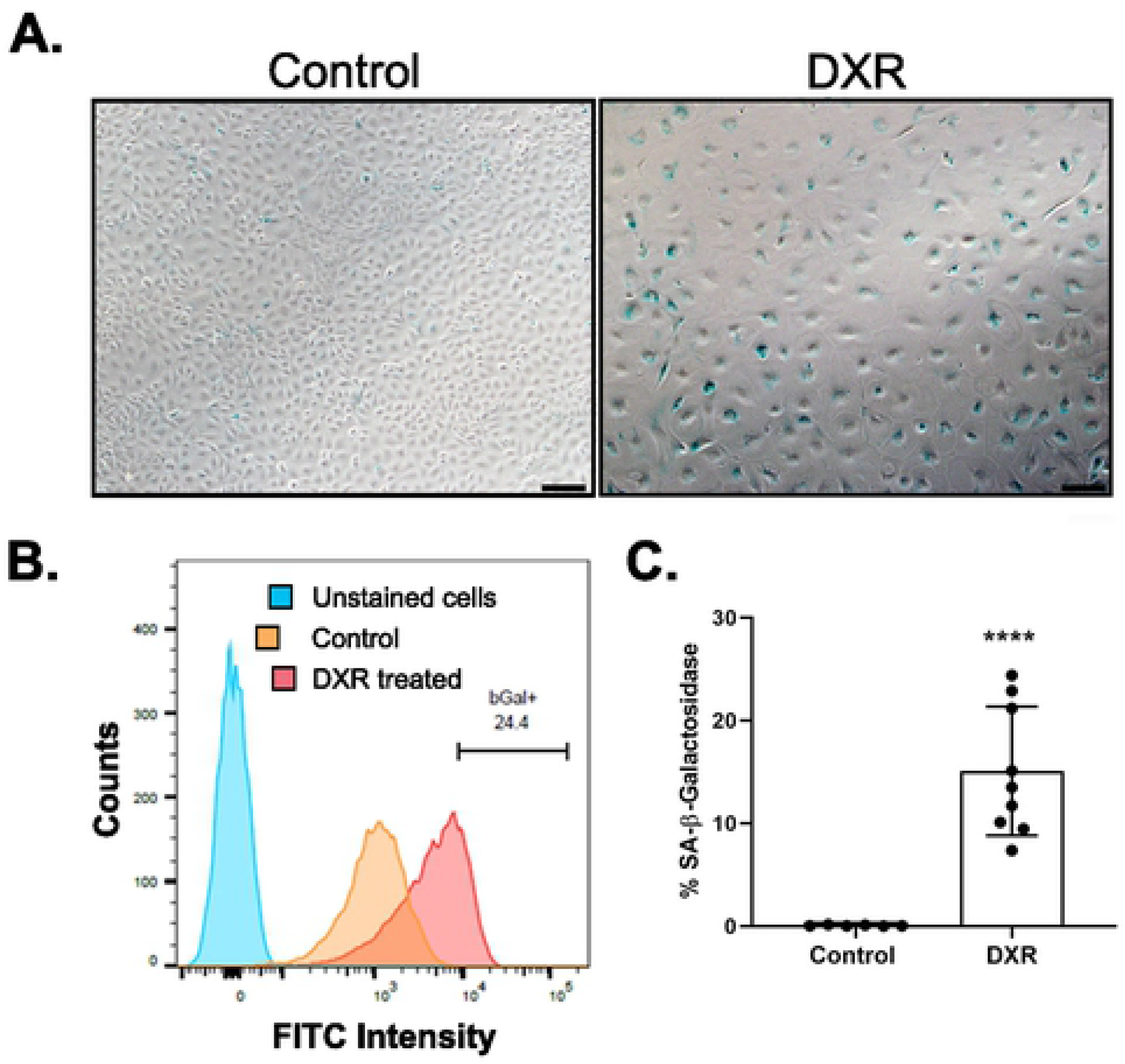
DXR treatment increase cell size. IEC-6 cells were cultured in the presence of vehicle or 0.2 μg/ml DXR for 4 days and evaluated for changes in cells size. (A) Representative flow cytometric analysis of forward and side scatter of IEC-6 cells after 4 days of vehicle (Control) or 0.2 μg/mL DXR (DXR). (B) Quantitative analysis of forward scatter (FSC) (n=6-9), error bars represent the standard deviation (SD). Significance was determined by unpaired Student’s t-test; **** P< 0.0001. (C) Representative brightfield micrographs of IEC-6 cells after 4 days of vehicle (Control) or 0.2 μg/mL DXR (DXR) treatment. (D) Superplot of cell area (μm^2^) comparing control and DXR treated IEC-6 cells. For statistical analysis n=6. Significance was determined by unpaired Student’s t-test; **** P< 0.0001. Scale bar equals 100 μm.

### DXR-treatment induces senescence associated cell senescence

Senescence associated-β galactosidase (SA-β-gal) activity is a hallmark of cellular senescence. IEC-cells were treated with vehicle or 0.2 μg/ml DXR for 4 days prior to analysis of SA-β-gal activity by colorimetric staining. As shown in Figure 4A, there were very few SA-β-gal positive cells in the vehicle treated control group but SA-β-gal activity was present in DXR treated cells. We next evaluated the proportion of SA-β-gal positive cells following vehicle or DXR treatment by flow cytometry using a green fluorogenic substrate for SA-β-gal. As seen in Figure 4B, when evaluated in the FITC channel, DXR treatment shifts IEC-6 cells to higher levels of FITC fluorescence intensity indicative of increased SA-β-gal activity. This is quantified in Figure 4C where we demonstrate a significant increase in the percentage of cells positive for SA-β-gal activity. Overall, SA-β-gal cellular staining confirmed that a concentration of 0.2 μg/mL of DXR is sufficient to induce senescence in IEC-6 cells.

**Figure 4.**
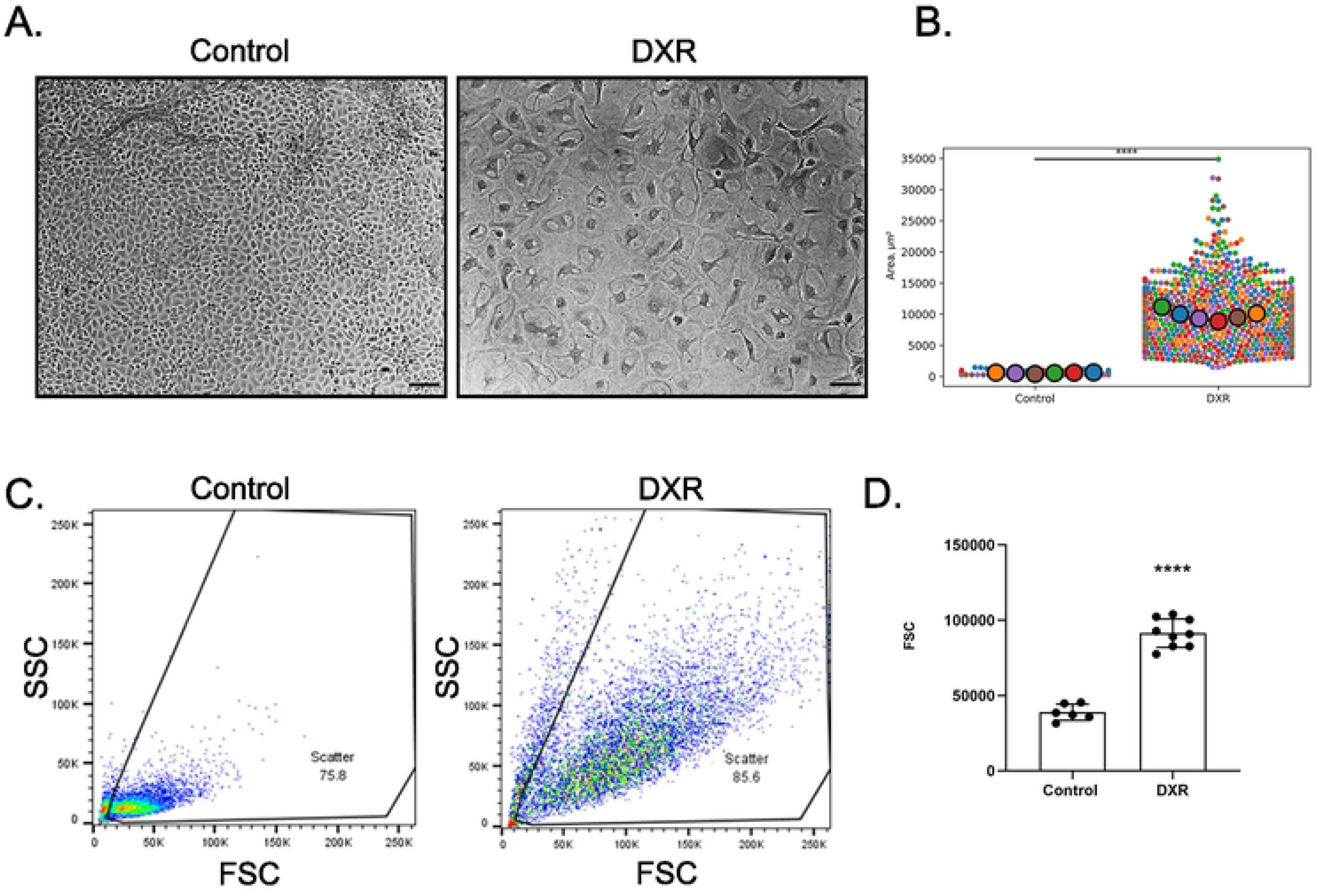
DXR treatment induces SA-β-Galactosidase activity. IEC-6 cells were cultured in the presence of vehicle (Control) or 0.2 μg/ml DXR (DXR) for 4 days and evaluated for senescence associated β-galactosidase activity (SA-β-Gal). (A) Representative images of IEC-6 cells demonstrating SA-β-Gal activity as assayed by the accumulation of intracellular blue precipitate. This blue precipitate is present in DXR treated IEC-6 cells (DXR) but is absent in vehicle treated cells (Control). Scale bar is equal to 100 μm. (B) Representative flow cytometry plot of SA-β-Gal activity in unstained (blue), Control (orange), and DXR treated (red) cells as assessed by intensity of accumulated green fluorescent enzymatic product. (C) Quantification of the proportions of SA-β-Gal positive vehicle (Control) and DXR-treated (DXR) cells (n=6-9), error bars represent the standard deviation (SD) from three different cell passages. Significance was determined by unpaired Student’s t-test; **** P< 0.0001.

### DXR induces genes of the senescence-associated secretory phenotype

Senescent cells express a cadre of genes including, *Il6*, *Cxcl1*, *Faim2*, *Ptges, and Cdkn1a*, as part of the Senescence-Associated Secretory Phenotype (SASP). Evaluation of mRNA expression of *Il-6*, *Faim2, Cdkn1a* and *Ptges* revealed differences between untreated control and DXR groups (Fig. 5). *IL6*, the most prominent cytokine of the SASP, exhibited a 12.91-fold increase in DXR treated cells (Fig. 5A). Senescence associated inflammatory responses, *Ptges* and *Faim2,* were also significantly upregulated (Fig. 5B and C), while *Cxcl1* (Fig. 5D) was not changed. *Cdkn1a*, which encodes the protein, p21, and is associated with cell cycle arrest, exhibited a 20.89-fold increase in DXR treated cells (Fig. 5E). Expression of *Sirt1*, which encodes the NAD-dependent deacetylase protein SIRT1 was decreased following DXR treatment (Fig. 5F).

**Figure 5.**
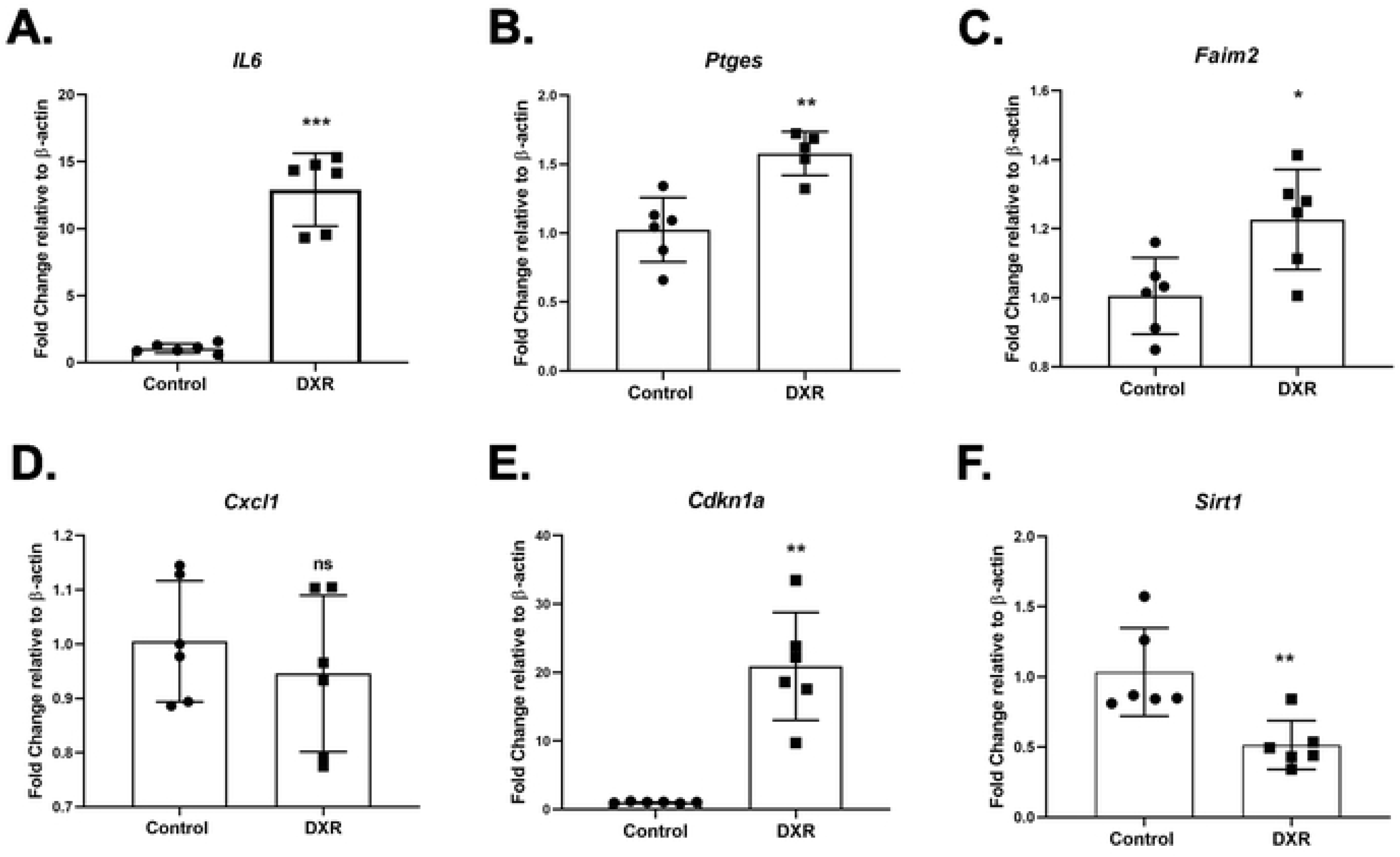
DXR treatment increases transcript levels of Senescence-Associated Secretory Phenotype genes. IEC-6 cells were cultured in the presence of vehicle (Control) or 0.2 μg/ml DXR (DXR) for 4 days and then lysed for isolation of total RNA. Total mRNA was subjected to qRT-PCR to quantify expression of senescence-associated secretory phenotype genes: (A) *IL-6*, (B) *Ptges*, (C) *Faim2*, (D) *Cxcl1*, (E) *Cdkn1a*, and (F) *Sirt1*. n=5-6 per group, error bars represent the standard deviation (SD). Significance was calculated using unpaired Student’s t-test, ns=not significant, * P<0.05, ** P < 0.01, *** P< 0.001.

### DXR-induced senescence is associated with changes in genes associated with intestinal stem cells and Wnt signaling

Silencing of intestinal stem cell and Wnt-signaling related genes has been observed in aging-associated senescence of intestinal epithelial cells. Therefore, we evaluated the expression of several candidate genes following treatment with DXR. Lgr5 expression was significantly decreased following DXR treatment (Fig. 6A), while Ascl2 expression was significantly increased (Fig. 6B). We found that Wnt-associated gene changed as well following DXR treatment. Axin2 was significantly decreased (Fig. 6C), Myc was significantly increased (Fig. 6D), and Rnf43 did not change (Fig. 6E).

**Figure 6.**
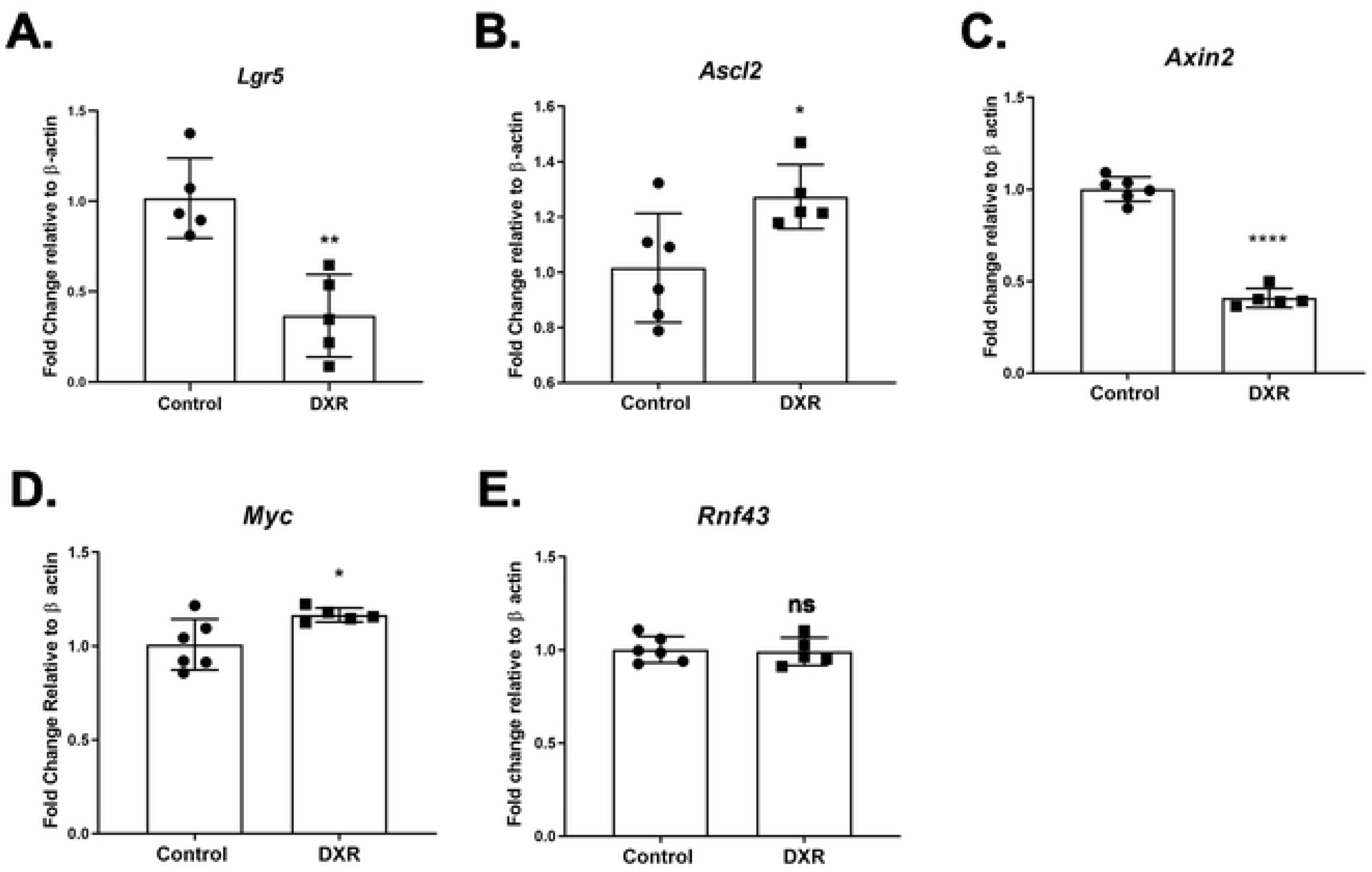
DXR treatment alters expression of genes associated with active intestinal stem cells and Wnt signaling. IEC-6 cells were cultured in the presence of vehicle (Control) or 0.2 μg/ml DXR (DXR) for 4 days and then lysed for isolation of total RNA. Total RNA was subjected to qRT-PCR to quantify expression of genes associated with active intestinal stem cells: (A) *Lgr5*, (B) *Ascl2*, (C) *Axin2*, (D) *Myc*, and (E) *Rnf43*. n=5-6 per group, error bars represent the standard deviation (SD). Significance was calculated using unpaired Student’s t-test, ns=not significant, * P < 0.05, ** P < 0.01, **** P < 0.0001.

### DXR reduced level of intracellular HMGB1 protein

Release of HMGB1, a known “alarmin” into the extracellular space as a result of cellular damage following DXR treatment has been demonstrated to activate immune cells. We therefore investigated whether HMGB1 is released from IEC-6 cells after DXR induced death and damage. IEC-6 cells were treated for 4 days with DXR and whole cell lysates were analyzed for intracellular levels of HMGB1 post DXR treatments via western blotting (Fig. 7A). DXR treatment significantly reduced the intracellular levels of HMGB1 compared with the untreated group (Fig. 7B), suggesting that DXR treatment decreases dramatically the level of intracellular HMGB1 protein in IEC-6 cells. Furthermore, analysis of *Hmgb1* mRNA expression (Fig. 7C) showed no significant differences between vehicle treated and DXR treated IEC-6 cells suggesting that the decrease in intracellular HMGB1 is due to its release from cells rather than from decreased expression.

**Figure 7.**
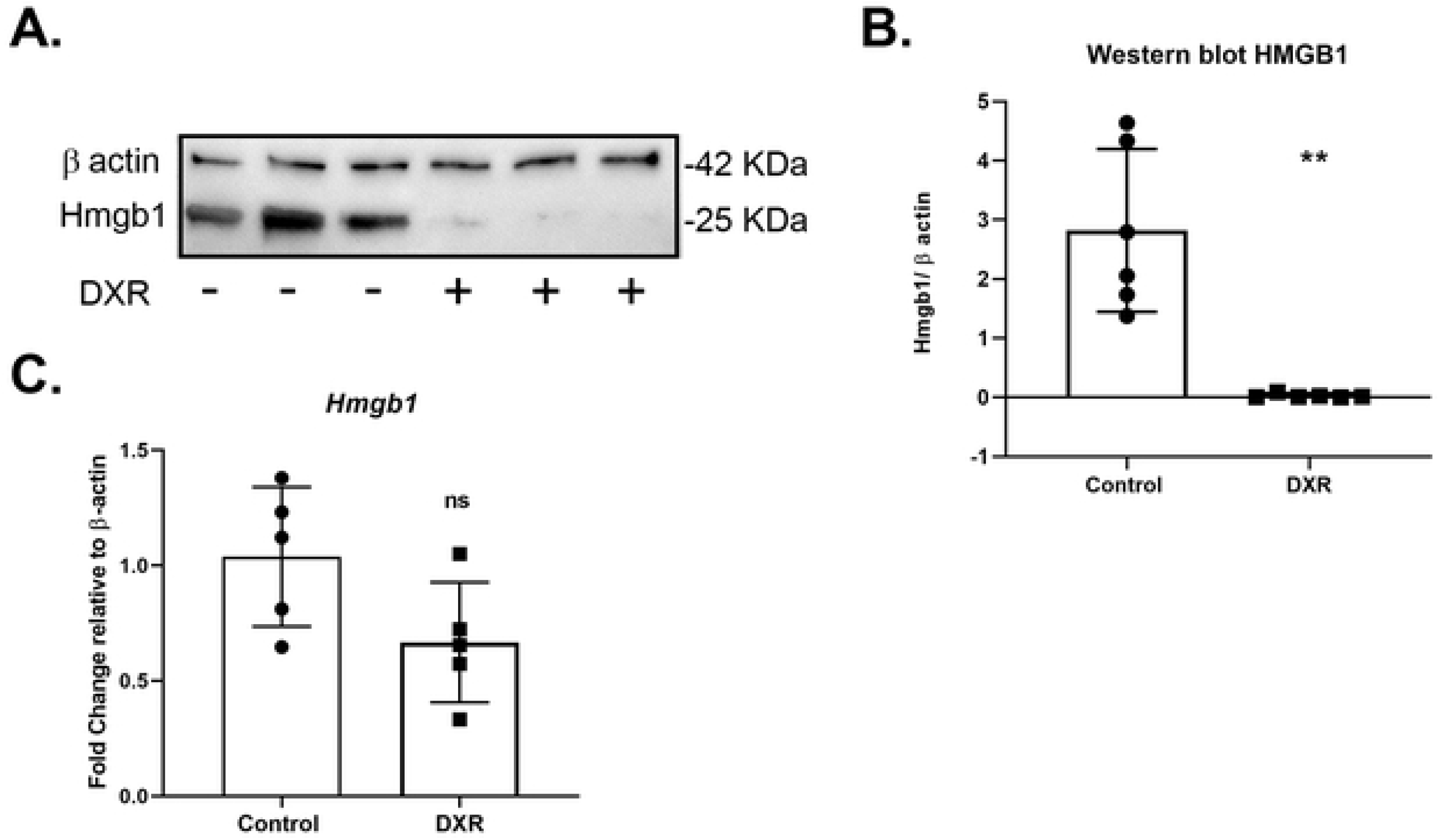
Effect of DXR on HMGB1 levels in IEC-6 cells. IEC-6 cells were cultured in the presence of vehicle (Control) or 0.2 μg/ml DXR (DXR) for 4 days and lysed for collection of total protein and total RNA. (A) Representative immunoblot of intracellular Hmgb1 protein and β-actin from IEC-6 cells treated with vehicle (−) or DXR (+). (B) Quantification of Hmgb1 protein expression relative to β-actin protein expression in Control and DXR-treated IEC-6 cells (n=6), **P<0.01. (C) mRNA expression of *Hmgb1* in Control and DXR-treated IEC-6 cells (n=5), ns=not significant. Significance was calculated using unpaired Student’s t-test.

## Discussion

Doxorubicin induces cellular senescence in numerous cell types as part of its mechanism of action.(8–11) In these cells, DXR-induced senescence also includes increased genomic instability, induction of autophagy, and upregulated expression of microRNAs.(8,12) However, whether DXR would induce senescence in intestinal epithelial cells was unknown. In this study, we demonstrated in cultured intestinal epithelial cells that DXR rapidly induces activation of cleaved caspase 3 in a subpopulation of cells, while in others it induces a senescent phenotype. This suggests a hierarchy of cellular sensitivity to DNA damaging agents similar to what we observe *in vivo* exists in culture. Induction of senescence by DXR in intestinal epithelial cells manifests as increased cell size, upregulation of SASP related genes, altered expression of gene associated with stemness, and decreased expression of SIRT1.

Data from this study are reminiscent of those from studies of intestinal stem cells from aged mice which demonstrate that senescent stem cells accumulate in intestinal crypts as mice age.(6,13) Accumulation of senescent cells is thought to contribute to decreased functional capacity, dysregulation of proliferation, and altered regenerative ability of ISC.(14,15) This leads us to consider that senescent cells could accumulate in small intestinal crypts following treatment with genotoxic agents, such as doxorubicin. Premature accumulation of such cells because of chemo/radiation therapy could account for altered intestinal function and could hasten potential impacts of age-associated senescence on function of the small intestine in the long term. Furthermore, similar to aging intestinal stem cells, DXR-induced senescence is accompanied by promoter hypermethylation and other epigenetic modifications that can alter expression of genes needed for stemness and proliferation.(16)

Our data show that senescent intestinal epithelial cells in culture release HMGB1 as is evidenced from the decrease in intracellular HMGB1 protein with no changes in HMGB1 mRNA expression. In senescent cells HMGB1, a conserved nuclear protein, is moved from the nucleus to the extracellular space where it serves as an alarmin.(17,18) Alarmins are detected by and activate cells of the immune system by binding to receptors such as Toll-like receptor 4 (TLR4) and receptor of advanced glycation end products (RAGE).(19–21) HMGB1 participates in DXR-induced damage of cardiomyocytes, and loss of TLR4 reduces DXR-induced intestinal damage.(22,23) TLR4 is expressed on numerous cell of the innate immune system, including macrophages and neutrophils, and is also expressed by active intestinal stem cells.(24–27) Binding of released HMGB1 to TLR4 on innate immune cells causes their activation, immune surveillance and elimination of senescent cells .(28–30) We have previously demonstrated that treatment of mice with DXR induces an acute inflammatory response that included infiltration of macrophages and neutrophils into the peri-cryptal space concomitant with crypt loss and mucosal damage.(31) The absence of enteric bacteria in germ free mice or pretreatment of mice with high-dose oral antibiotics to deplete enteric bacteria prevented this infiltration of immune cells and ameliorated crypt loss.(31,32) This correlation between the bacteria, innate immune cells and crypt loss could involve the interaction of epithelial cell-derived alarmins with infiltrating immune cells and should be further studied.

Finally, our results suggest that proliferative intestinal epithelial cells can have disparate responses to exposure to DXR by either becoming apoptotic or senescent. While it is known that p53 mediates both the apoptotic (via PUMA) and the survival (via p21) pathways, the molecular triggers within a cell that dictate which pathway is taken are less well understood. Currently, our understanding is that *in vivo* DXR preferentially induces apoptosis in highly proliferative epithelial cells of the small intestinal crypt, including the active intestinal stem cells, and that within the remaining crypt epithelium are cells capable of resisting DNA damage and regenerating the depleted epithelium. That a cohort of these remaining cells could be senescent adds another potential layer of complexity to the cellular dynamics of intestinal epithelial repair. Whether there is evidence of induction of senescence in these cells, the impact these cells have on crypt epithelial cell function, and the fate of these cells warrants further study.

## Acknowledgements

Flow Cytometry experiments were performed in the Flow Cytometry and Cell Sorting facility at North Carolina State University-College of Veterinary Medicine.

